# A Recurrent Neural Network Model for Flexible and Adaptive Decision Making based on Sequence Learning

**DOI:** 10.1101/555862

**Authors:** Zhewei Zhang, Huzi Cheng, Tianming Yang

## Abstract

The brain makes flexible and adaptive responses in the complicated and ever-changing environment for the organism’s survival. To achieve this, the brain needs to choose appropriate actions flexibly in response to sensory inputs. Moreover, the brain also has to understand how its actions affect future sensory inputs and what reward outcomes should be expected, and adapts its behavior based on the actual outcomes. A modeling approach that takes into account of the combined contingencies between sensory inputs, actions, and reward outcomes may be the key to understanding the underlying neural computation. Here, we train a recurrent neural network model based on sequence learning to predict future events based on the past event sequences that combine sensory, action, and reward events. We use four exemplary tasks that have been used in previous animal and human experiments to study different aspects of decision making and learning. We first show that the model reproduces the animals’ choice and reaction time pattern in a probabilistic reasoning task, and its units’ activities mimics the classical findings of the ramping pattern of the parietal neurons that reflects the evidence accumulation process during decision making. We further demonstrate that the model carries out Bayesian inference and may support meta-cognition such as confidence with additional tasks. Finally, we show how the network model achieves adaptive behavior with an approach distinct from reinforcement learning. Our work pieces together many experimental findings in decision making and reinforcement learning and provides a unified framework for the flexible and adaptive behavior of the brain.

## Introduction

Consider a scenario in which a cheetah sneaks up on a deer until it starts the final dash to catch it. Every move has to be calculated carefully based on the sensory inputs, for example, the distance to the deer and the surrounding environment, and the past experience, for example, the deer’s speed. Furthermore, the cheetah should be able to predict how the deer would respond to its move to allow more timely adjustment of its actions. This is an example of a central function of the brain: to generate responses to maximize the gain based on a continuous stream of sensory inputs and movements that reflect a complicated and volatile world. The contingencies that the brains need to learn should take into account the sensory inputs, the organism’s own actions, and the resulting reward outcomes, all strung together into a sequence.

A variety of theoretic models have been used to investigate certain components of this general contingency problem. For example, in the perceptual decision-making and sensory-motor transformation field, the research question often centers around how sensory information as evidence is integrated and leads to specific actions. Theoretical models such as the drift-diffusion model (Ratcliff, 1978; Stone, 1960) and attractor neural network models (X.-J. Wang, 2001; Wong & Wang, 2006) have gained a lot of success in both modeling the behavior and explaining the neuronal response patterns in the brain. Yet these studies often do not take into account the volatile nature of the environment. In contrast, the conditioning and learning field focus on the learning of the associations between reward or punishment outcomes and other events in a volatile environment. The reinforcement learning (RL) framework has become a standard approach to understand the behavior and the brain circuitry in this field (Dayan & Niv, 2008; Schultz, Dayan, & Montague, 1997; Sutton & Barto, 2012). In RL, state and action values are updated when reward feedbacks differ from expectations. The framework provides an explanation on how animals gradually learn the values associated with stimuli and actions and select actions accordingly. Yet, RL does not usually deal with the problem of inferring states based on noisy sensory inputs. It is inefficient for learning complicated decision-making tasks, especially those with a large state space. More problematic is that in most studies RL assumes that states are observable and task structures are known, which is not true in the real world. Biologically realistic neural network models of RL that acquires task structures through learning have been proposed but they are limited to rather simple structures (J. X. Wang et al., 2018; Zhang, Cheng, Lin, Nie, & Yang, 2018).

So far, there is a lack of a model general or flexible enough to cover different aspects of decision making and learning. A more general framework is necessary for furthering our understanding of the brain. Here, we propose a neural network model based on sequence learning to model the flexible and adaptive behavior and to understand the underlying neural mechanisms. The network takes inputs in the form of event sequences and is trained to make predictions of future events with supervised learning. Importantly, the input event sequences include not only the sensory events, but also the action events and the reward outcome events, which allow the network to establish the contingencies between all three types of events. The outputs of the network serve as a prediction of future events and the network is trained to minimize the difference between the predictions and the future input sequences. The predictions of action events are used to generate the model’s actual responses, which are assessed for the network’s behavior performance. The predictions of sensory and reward events are not directly exhibited in behavior, but they are part of the learning and have important implications in modeling the brain.

To demonstrate how the network model works, we choose four exemplary behavior tasks that have been previously used in animal and human studies, each covering a different aspect of decision making and learning. The network is not constructed for any particular task. Instead, it learns the task structure and task logic through training. The first task, a reaction time version of probabilistic reasoning task, is one of the most complicated tasks in the literature that provide us both behavior and single unit recording data to evaluate the model (Kira, Yang, & Shadlen, 2015; Yang & Shadlen, 2007). We use it to demonstrate the capability of the network to learn complex contingencies between events in long sequences and the similarity of the units in the network to the previous neurophysiological findings. We further use a multi-sensory integration task to show how the network generalizes Bayesian inference across different sensory modalities (Gu, Angelaki, & DeAngelis, 2008). The third task, a post-decision wagering task, is chosen to show that the model can support meta-cognition such as confidence (Kiani & Shadlen, 2009). Last but not least, the final task is adapted from the two-step task, first described to illustrate model-based RL (Daw, Gershman, Seymour, Dayan, & Dolan, 2011). The task requires the learning to be extended across trials, which allows us to demonstrate how the network accounts for adaptive behavior with a distinct approach from RL.

Together, our results show that a network model that is simply trained to make predictions based on event sequences may account for a large body of experimental findings from both the decision making and the reinforcement learning field. Not only does the model reproduce the animals’ and humans’ behavior in previous studies, the units in the network also show response patterns resembling those of the neurons in the brain. The model makes testable hypotheses on the neuronal response patterns and circuit structures in the brain and suggests novel interpretations for some of the previous experimental work. These results suggest the potential of our model to be a unified framework for decision making and learning that may reveal the computational principle in the brain.

## Results

### Network

Our network model contains three layers: the input layer, the hidden layer based on gated recurrent units (GRU), and the output layer (Fig 1a). The input layer contains units that carry the information about the sensory, motor, and reward events on a timeline. Each unit’s activity can be a binary variable, representing the presence or the absence of the corresponding event, or a continuous variable representing stimulus strength. The input units are fully connected with the next layer. We use vector *x_t_* of length *N_IL_* to describe the activities of the input layer units.

**Figure 1.**
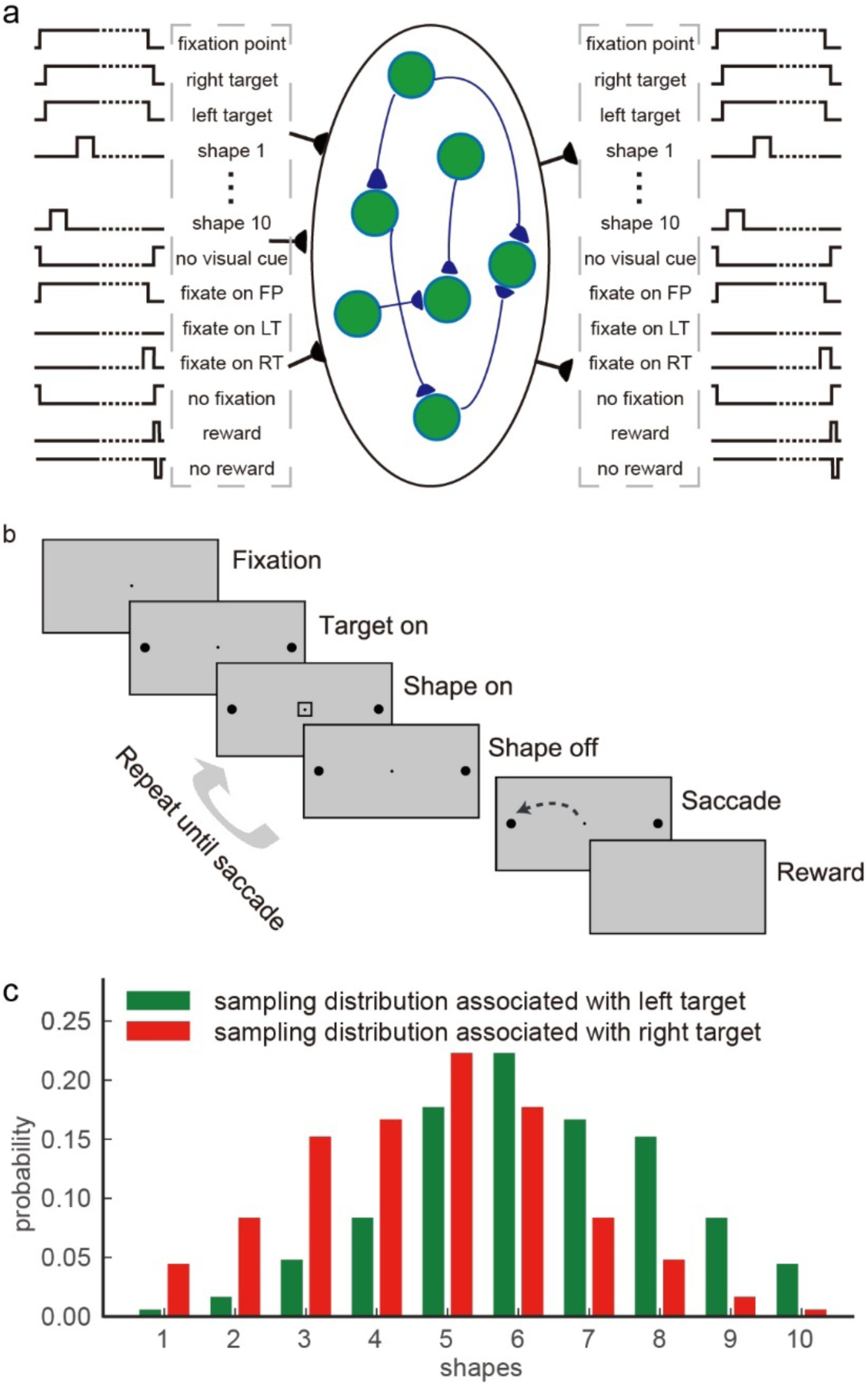
**a**. The model diagram. The network has three layers: the input layer, the gated recurrent unit layer, and the output layer. The input layer receives the input sequences of sensory events, action events, and reward events. The GRU layer has 128 gated recurrent units. The output layer units mirror the input layer units and represent the prediction of the future events. The diagram illustrates the particular input and output units for the Task 1. **b**. Task 1: the reaction-time version of probabilistic reasoning task. The subject fixates at a central point and views a sequence of shapes to make a decision by moving the eyes toward one of the two choice targets in the peripheral. Each shape confers information regarding to which target will be rewarded. An optimal strategy is to integrate the information and make a choice when the integrated information hits a bound. **c**. The sampling distributions. Shapes are sampled from the sampling distribution associated with the correct target in each trial.

The hidden layer is the core of our model. It is adapted from the GRU network, which has been shown to be suitable for learning sequences (Cho et al., 2014; Chung, Gulcehre, Cho, & Bengio, 2014). There are *N_h_*=128 units in the hidden layer. Each gated recurrent unit’s activity is a nonlinear combination of both the inputs and its activity at the previous time step, which are regulated by an update gate and a reset gate. The nonlinear reset gate has been shown to be critical for the network to learn long contingencies across long temporal sequences (Greff, Srivastava, & Koutník, 2016). The state vector *h_t_* of length *N_h_* describes the responses of these units at time *t*, which is updated with the information from the input layer as follows:

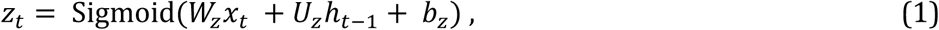

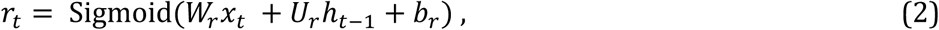

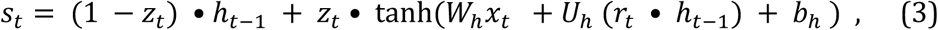

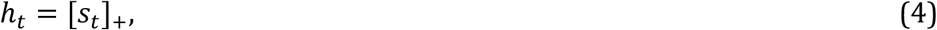

where *z_t_* and *r_t_* represent the update gate vector and the reset gate vector, respectively; *W_z_*, *W_r_* and *W_h_* are the input connection weight matrices for the update gates, the reset gates, and the gated units; *U_z_*, *U_r_* and *U_h_* are the recurrent connection weight matrices for the update gates, the reset gates, and the gated units; *b_z_*, *b_r_* and *b_h_* are the bias vectors for the update gates, the reset gates, and the gated units; and • indicates the element-wise multiplication. The rectified-linear function [•]_+_ keeps the responses *h_t_* non-negative. The initial value of the state vector *h_t_* at the beginning of each trial is reset to zero.

The hidden layer units project to the output layer with full connections. The output layer is composed of an array of units that mirror the input layer units, representing the network’s predictions of the corresponding sensory events, action events and reward events for the next time step. The activity of the output layer units *o_t_* (a vector of length *N_OL_*) is:

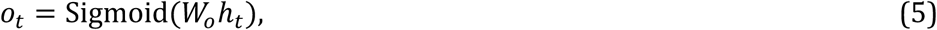

where *W_o_* is the output connection weight matrix. The prediction y is a function of response *o_t_*:

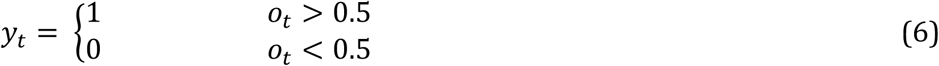

The network predicts the corresponding event would happen in the next time step when 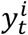 is 1. If 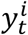 corresponds to an action, the probability of the action being carried out at the next time step is a softmax function of *o_t_*:

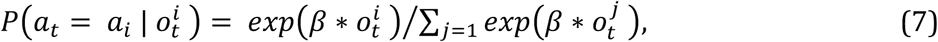

where *β* is the temperature. The chosen action is evaluated for the network’s task performance for testing purposes.

Fig 1a illustrates the network structure with the inputs and outputs corresponding to the Task 1 that we discuss below. For the other tasks, it is only necessary to change the input and output units to reflect the relevant events.

The goal of the training is for the network to learn to predict events and generate sequences that lead to a reward. The loss function *L* is defined as the sum of mean squared error between elements in the output *o_t_* and actual event sequence *e_t_*_+1_ for all time points *t*.

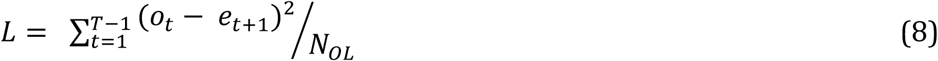

where *T* is the length of each trial. The parameter vector *θ* (including *W*_*_, *U*_*_ and *b*_*_) of the network is updated with the Adam (Kingma & Ba, 2014) and the gradient clipping is applied to avoid exploding gradients as follows:

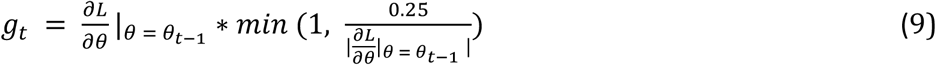

### Task 1: Probabilistic reasoning task

We train our model with a reaction-time version of the probabilistic reasoning task that was used to study the neural mechanism of sequential decision making (Kira et al., 2015). The task is illustrated in Fig 1b. In this task, a subject has to make decisions between a pair of eye movement targets. Initially, the subject needs to fixate on a central point on a computer screen. Then a stream of shapes appears sequentially near the fixation point. There are totally 10 possible shapes. Each shape conveys information on the correct answer. The subject needs to integrate information of the shapes to form a decision and move the eyes to look at the choice target whenever ready.

The contingency between the shapes and the targets is described by two distributions. Each target is associated with a distribution of the appearance probability of the shapes in the sequence (Fig 1c). In each trial, the computer randomly picks the correct target. Each shape in the sequence is independently sampled from the distribution associated with the correct target. Because the likelihoods of observing a particular shape under the two distributions are different, each shape provides information on which target is correct. It has been shown that the sequential probability ratio test (SPRT) is an optimal strategy to solve the task in the sense that it requires the least number of observations to achieve a desired performance (Wald & Wolfowitz, 1948). In the SPRT, one needs to accumulate the log likelihood ratio (logLR) associated with each piece of evidence, which is the log ratio between the conditional probabilities of observing the evidence given the two testing hypotheses. In our task, the logLRs associated with each shape range from -0.9 to 0.9 (base 10). We define positive and negative logLRs as evidence supporting the left and the right target, respectively.

The task has several attractive features for our modeling purposes. First of all, the task features a sophisticate statistical structure containing the shapes, the choice, and the reward. As the shapes appear one by one, an ideal observer not only gains information on what would be the appropriate choice, but also can deduce how likely a particular shape will appear next and how likely a reward can be expected. Second, this is a reaction time task in which choices have to be made at appropriate time to achieve a certain tradeoff between speed and accuracy. This allows us to demonstrate the flexibility of our model for learning sequences of variable length. Last but not least, the task is one of the most complicated tasks that have been used in animal studies with both behavior and neural data available. We not only can compare the behavior between the model and the animals, but also can look into the network and study the network units’ activities and compare them against the experimental findings.

### The training dataset

We train the network with task event sequences created with simulated trials in which choices are generated with a drift-diffusion model with collapsing bounds. In each trial, a correct target is randomly determined, and the shapes are generated with its associated distribution (Fig 1c). A choice is triggered when the accumulated logLR reaches to either of the two opposite collapsing bounds. The bounds start at ±1.5 and linearly collapse toward 0 at the rate of 0.1 per shape epoch. The left target is chosen if the positive bound is reached, and the right target chosen if the negative bound is reached. If the choice matches the pre-determined correct target, a reward is given, and the corresponding sequence is included in the training dataset.

The results presented below are based on 20 simulation runs using a training set generated with 7.5 × 10^5^ simulated trial sequences.

### Behavior Performance

After training, the network performs the task well. When the total logLR associated with the shape sequence is positive, the network tends to choose the left target, and when the total logLR is negative, it tends to choose the right target (Fig 2a). A logistic regression reveals that the logLR assigned to each shape correlates well with its leverage on the choice (Fig 2b). The mean reaction time, quantified as the number of shapes that the model uses for decision, is 6.36±0.04 (mean±s.e.m.). Both the distribution of the reaction time and the total logLR at the time of choice suggest that the model behavior is consistent with the DDM with collapsing bounds (Fig 2c). We use another logistic regression to examine the effects of shape order on the choice. The first shapes in the sequence exert similar leverages on the choice to the rest of the shapes, except for the last shape, suggesting all shapes except the last are used equally in decision making (Fig 2d). The last shape’s sign is almost always consistent with the choice (> 99% of all trials). This is also consistent with the DDM, in which the last shape brings the total logLR over the bound. Overall, the network performance resembles, although understandably better than, the behavior of the macaque monkeys trained with a very similar task (Kira et al., 2015).

**Figure 2.**
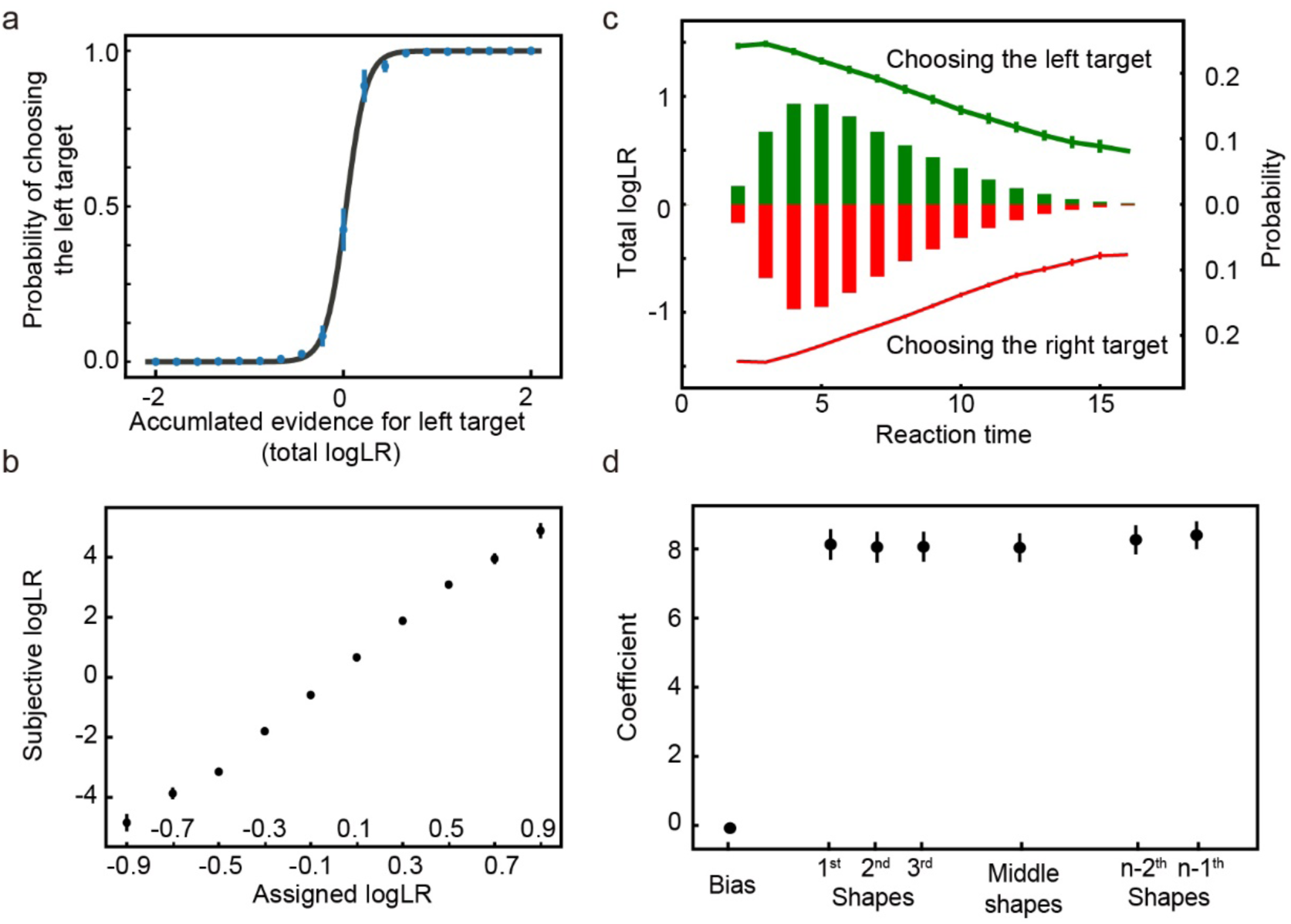
**a**. The psychometric curve. The model more often chooses the target supported by the accumulated evidence. The black curve is the fitting curve from the logistic regression. **b**. The leverage of each shape on choice revealed by the logistic regression is consistent with its assigned logLR. **c**. Reaction time. The bars show the distribution of reaction time, quantified by the number of observed shapes (right y-axis). Green and red indicate the left and right choices, respectively. The lines indicate the mean total logLR (left y-axis) at the time of decision, grouped by reaction time. Trials with only 1 shape or more than 16 shapes comprise less than 0.1% of the total trials and are excluded from the plot. **d**. The leverage of the first 3, the second and third from the last, and the middle shapes on the choice. Only trials with more than 6 shapes are included in the analysis. No significant differences are found between any pair of the coefficients of shape regressors (two-tailed t test with Bonferroni correction). The error bars in all panels indicate S.E. across runs. Some error bars are smaller than the data points and not visible.

The good performance of the network is not because the trials sequences in the testing dataset overlap with those in the training dataset significantly, which is not true because the possible number of trial sequences in this task is astronomical and each shape sequence in the training and testing dataset is randomly generated. The network is able to generalize beyond the training dataset. With a training dataset that contains only 1000 unique trial sequences, the network can still achieve a good performance (Supplementary Figure 1). Therefore, the learning of the statistical structure of the task is likely the reason for the model’s performance.

### Network analyses – Evidence and Choice encoding

Next, we examine how the evidence and the choice are encoded in the network. The example unit in Fig 3a shows a classical ramping-up activity pattern that has been reported in neurons from the prefrontal cortex (Kim & Shadlen, 1999), the parietal cortex (Kira et al., 2015; Roitman & Shadlen, 2002; Yang & Shadlen, 2007), and the striatum (Ding & Gold, 2010). Its activity increases when the total logLR grows to support its preferred choice and decreases when the total logLR is against its preferred choice. The responses converge around the time when its preferred choice is chosen. The population analysis using all the units that are selective to the total logLR finds the same pattern (Fig 3b).

**Figure 3.**
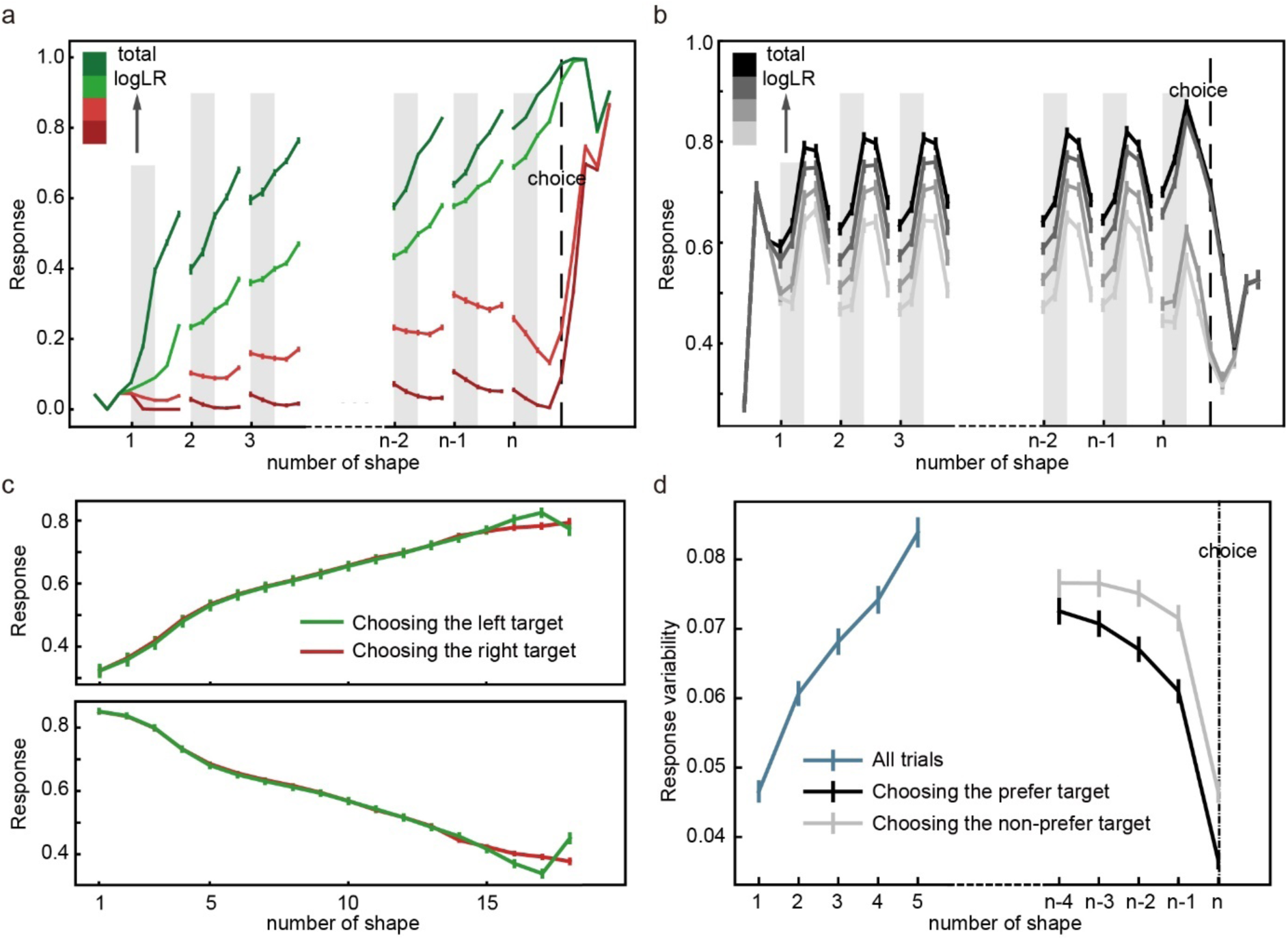
**a**. An example unit that prefers the left target. Its activity increases when the evidence supporting the left target grows and decreases when it drops. The unit’s responses converge when the network chooses its preferred target. The trials are grouped into quartiles by the accumulated evidence in each epoch, which are indicated with color. The error bars indicate the S.E. across trials. **b**. Population responses of the units that are selective to the total logLR. The trials are grouped based on the accumulated evidence supporting each unit’s preferred target in each epoch. The error bars in panels **b, c** and **d** indicate the S.E. across units. c. Urgency units. Their activities ramp up (upper panel) or down (lower panel) regardless of the choice. **d**. Network unit response variability. The neurons’ response variability increases initially (blue curve) but decreases before the choice, more so when the preferred target is chosen (black) than when the non-preferred target is chosen (grey). Only the trials with more than five shapes are included in panel **a, b** and **d**.

In addition to the units that accumulate evidence and reflect the decision making process, we also find units that show a ramping-up or ramping-down activity pattern independent of the choice (Fig 3c). Their activities indicate the passage of time and may be interpreted as an urgency signal. Neurons with a similar activity pattern have been reported in the global pallidus (Thura & Cisek, 2017).

We use a linear regression to quantify the selectivity of each units (see Methods) to three important parameters for this task: the accumulated evidence, quantified with the total logLR, the choice outcome, and the urgency. During the shape presentation period, a large portion of units is found to encode the accumulated evidence (N = 63.00 ± 2.49), the choice outcome (N = 50.60 ± 2.03) and the urgency (N= 86.50 ± 1.10).

Finally, we calculate the response variability of the units in the hidden layer to study the dynamics of the population responses of the network (Fig 3d). This is analogous to the variance CE that Churchland and colleagues studied previously in the LIP neurons (Churchland et al., 2011). Our analyses are more straightforward as the units in our model do not have intrinsic Poisson noise. The response variability here is simply calculated as the standard deviation of the units’ responses across trials. We find that the response variability increases initially (linear regression, k = 0.0088, p < 0.001) as more shapes are presented and the evidence is accumulated. However, when we align the trials to the choice, a different pattern emerges. In the trials in which the preferred target is chosen, the response variability decreases before the choice (linear regression, k = -0.0038, p < 0.001; Spearman’s rank correlation, r = -0.045, p < 0.01) and reaches the minimum around the time of choice. In contrast, in the trials in which the non-preferred target is chosen, the response variability is significantly higher and its decrease is not significant until the last shape (linear regression, k = -0.0017, p = 0.07; Spearman’s rank correlation, r = -0.017, p = 0.28). The overall pattern is very similar to that of the LIP neurons reported previously (Churchland et al., 2011).

### Network analyses – When and Which

The balance between the speed and accuracy is an important aspect of decision making. The control of the speed and accuracy balance has been suggested to be exerted through the same neurons that accumulate evidence (Ding & Gold, 2012; Hanks, Kiani, & Shadlen, 2014; Heitz & Schall, 2012) or a distinct populations of neurons that reflect only the speed but not the choice (Thura & Cisek, 2017). We analyze the units’ activities and connectivities in the model to find out which is the case here in our model.

The network has three output units related to the choice. They include a unit for the fixation (FP) and two units for the saccades toward the left (LT) and the right (RT) target, respectively. We examine the connection weights between the hidden layer unit and the output units to understand how hidden layer units drive the choices. For each hidden layer unit, we estimate how it contributes to the outputs by defining two indices, *I_when_* and *I_which_*:

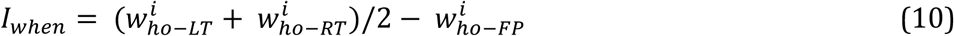

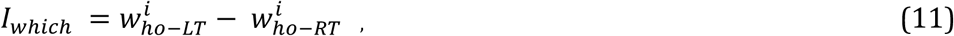

where 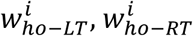 and 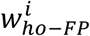 are the connection weights between the hidden layer unit *i* and the output layer units LT, RT and FP, respectively. Units with large positive *I_when_* promote the saccades over fixation, regardless of the direction, thus affect the reaction time. In contrast, units with large *I_which_* bias the saccade direction toward their preferred direction.

With these indices defined, we select two groups of hidden layer units: the *when* units and the *which* units. Within each group, we further divide them into a positive subgroup and a negative subgroup, depending on their signs of *I_when_* or *I_which_*. The +*when* units are selected with the following procedure. First, we sort all units with positive *I_when_* values by their *I_when_* values in the descending order. Then, we calculate the accumulative *I_when_*along this axis and select the top units that together contribute more than 50% of the sum of the positive *I_when_*. These units are defined as the +*when* units (10.70 ± 0.26 out of 50.60 ± 0.89 units with positive *I_when_*). They have larger connection weights to the two saccade output units than to the fixation output unit (Fig 4a). We use similar procedures to select the -*when*, the +*which*, and the -*which* units (n=13.80 ± 0.27 out of 77.40 ± 0.89 units with negative *I_when_*, 9.80 ± 0.30 out of 64.00 ± 1.25 units with positive *I_which_*, and 9.50 ± 0.28 out of 64.00 ± 1.25 units with negative *I_which_*, respectively). The -*when* units have smaller connection weights to the saccade units than to the fixation unit, the +*which* units have larger connection weights to the left saccade than to the right saccade output unit, and the -*which* units have smaller connection weights to the left saccade than to the right saccade output unit (Fig 4a). With these selection criteria, the *when* units are supposed to contribute more to the decision of when a response should be made, while the *which* units contribute more to the decision of which target should be chosen. Together, they constitute 34.22 ± 0.52% of all the units in the hidden layer.

**Figure 4.**
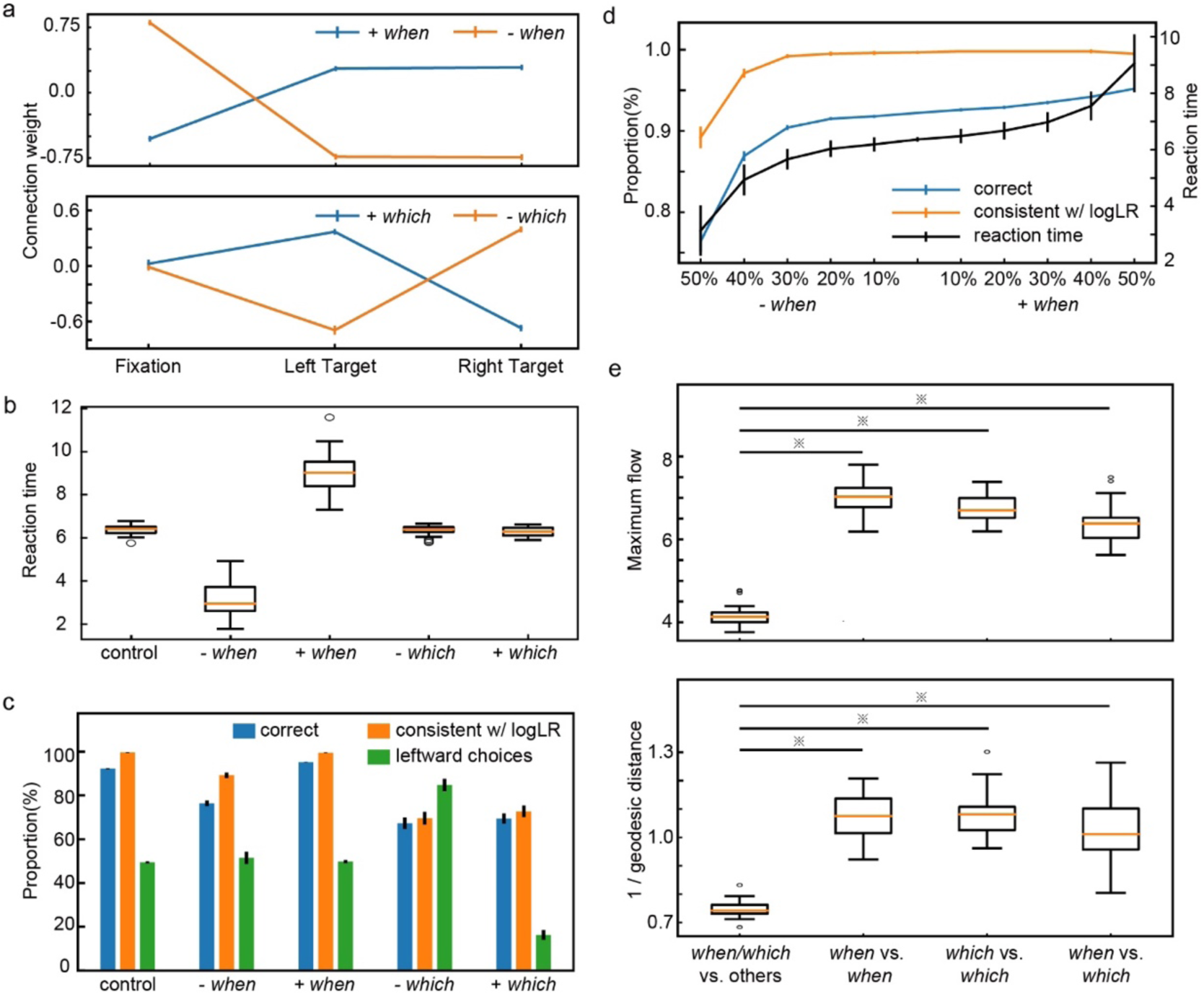
**a**. The connection weights between the eye movement output units and the *when* units (upper panel) and the *which* units (lower panel). **b**. The effect of lesions to the *when* and *which* units on the reaction time. **c**. The effect of lesions to the *when* and *which* units on the choice. The blue bars indicate the proportion of correct trials. The orange bars indicate the proportion of trials in which the choice is consistent with the sign of the accumulated evidence at the time of choice. The green bars indicate the percentage of trials in which the model chooses the left target. **d**. Speed-accuracy tradeoff. We suppress the output of a different portion of +*when*/-*when* units (see Methods). As more +*when* units’ outcome is suppressed, the model’s reaction time (black curve, right y-axis) increases along with the accuracy (blue curve, left y-axis). However, the proportion of trials in which the choices are consistent with the evidence (orange curve, left y-axis) stays the same except for the extreme cases. **e**. The maximum flow (upper panel) and the inverse of the geodesic distance (lower panel) between different unit groups. The smaller maximum flow and the larger geodesic distance between *when*/*which* units and other units suggest the relatively tight connections between the when and which units. ※ indicates significant difference (p<0.05, Two tailed t-test with Bonferroni correction). The error bars in all panels indicate the S.E. across runs.

Next, we examine how the ±*when* and ±*which* units affect the network’s behavior. The causality link can be demonstrated by simulated lesions that selectively inactivate the connections of each group of units to the output layer. With only the outputs inactivated, the hidden layer itself is not disturbed. When we inactivate the +*when* units, the RTs become longer while the accuracy remains intact (Fig 4bc). In contrast, when we inactivate the output of the -*when* units, the network’s RTs become smaller (Fig 4b). Although the network performance has an apparent drop, the network’s choice still accurately reflects the accumulated logLR, suggesting the performance drop is largely due to the fact that the network has to work with a smaller total logLR due to the shorter RT (Fig 4c). In comparison, when *which* units are manipulated, only the choice accuracy is affected but the RT remains the same (Fig 4bc). More specifically, inactivating +*which* units leads to a bias toward the right target, while inactivating - *which* units leads to a bias toward the left target (Fig 4c). These results suggest there are two distinct populations of hidden layer units contributing to the choice and the reaction time.

The way how *when* units affect the accuracy and reaction time suggests a possible mechanism to modulate the speed-accuracy tradeoff. We demonstrate this by suppressing different amount of *when* units’ outputs (see Methods for details). As more +*when* units’ output is suppressed, the reaction time becomes larger (Spearman’s rank correlation, p<0.001) and the accuracy becomes higher (Spearman’s rank correlation, p<0.001) (Fig 4d). Importantly, the choices are still consistent with the accumulated evidence except for the most extreme cases.

The distinct functional roles of the *when* and *which* units may suggest they have different connection patterns. We use the connection matrix of the hidden layer units to construct a weighted directed graph and calculate the geodesic distance and the maximum flow between the hidden layer units (see Methods). To account for the units that are not connected to each other, we use the average maximum flows and inverse of geodesic distance in the analysis, which is 0 for the unit pairs that are not connected.

Interestingly, we find the inverses of the geodesic distance between the *when* units and the *which* units are significantly larger than that between the *when*/*which* units and others. The maximum flows between the *when* units and the *which* units are also significantly higher than the network average (Fig 4e). These results suggest that the *when* and the *which* units belong to a tightly connected sub-network within the hidden layer and are not topologically separated.

### Network analyses – Predictive Coding

So far, we have shown that our network is able to generate appropriate choices based on shape sequences. However, the network is constructed and trained in the way that the sensory events are treated exactly the same as the action events. Therefore, the activities of the output units that correspond to the sensory events should provide predictions on the sensory events. In other words, our network may also serve as a generative model for the predictive coding framework.

In the probabilistic reasoning task, each choice is associated with its respective shape distribution. Therefore, as one accumulates information regarding to the choice, the probability distribution of the upcoming shapes can also be inferred. We plot the mean subthreshold activity, *o_t_*, of the ten shape output units in Figure 5a at the time point right before the onset of each shape. When the current evidence is in favor of the left target, the activities of the output layer shape units resemble the probabilities from which the shapes are drawn when the left target is the correct target. When the evidence is in favor of the right target, the activity pattern of the output layer shape units shifts and becomes more similar to the sampling distribution associated with the right target (Fig 5a).

**Figure 5.**
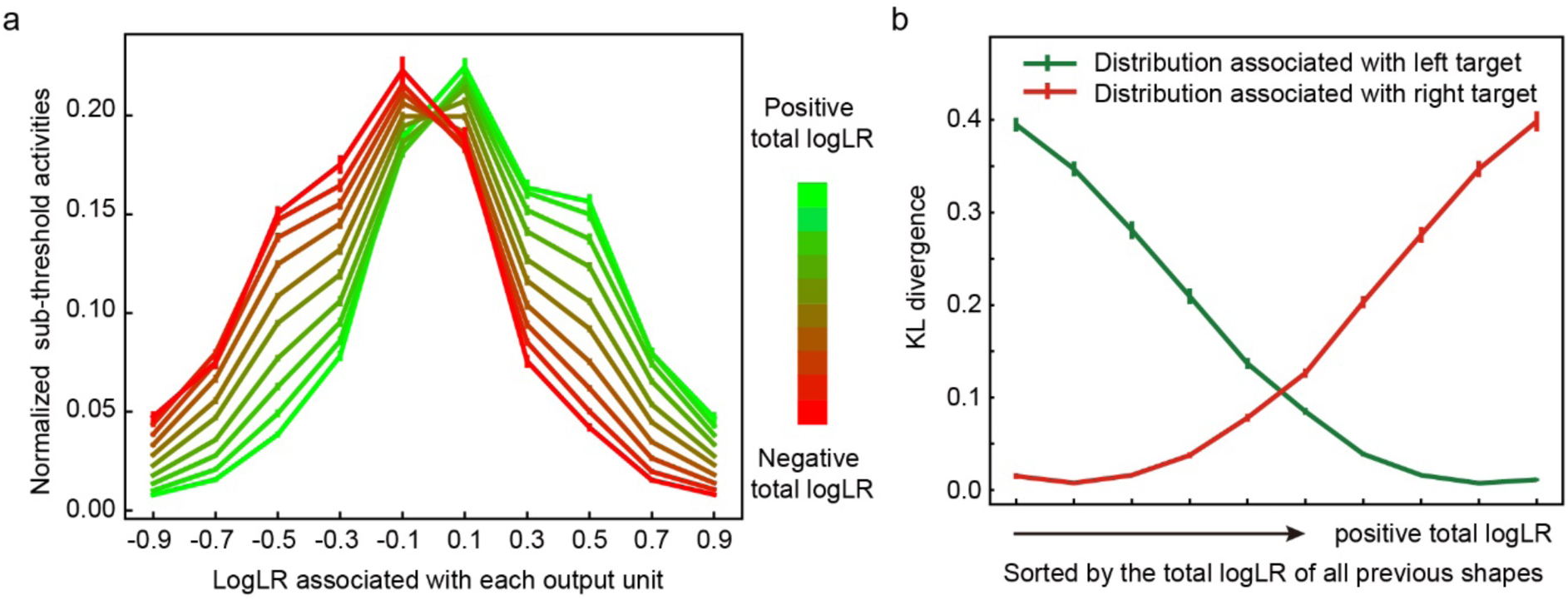
**a**. The normalized subthreshold activities of 10 shape output units. We calculate the 10 shape units’ activities at the time step immediately before each shape onset for all epochs in all trials in the test dataset (96555 trials and 583445 epochs from 20 runs), normalized by dividing each unit’s activity by the sum of activities of all 10 shape output units. Data are divided into 10 group by the total logLR before the shape onset, which is indicated by the color. **b**. The Kullback-Leibler (KL) divergence between the normalized subthreshold activities (as shown in the Fig. 5a) and the sampling distributions (shown in the Fig.1c). Data are grouped by the total logLR. The error bars indicate the S.E. across runs.

We quantify the similarity between the activity patterns of the output layer shape units and the sampling distributions by calculating the Kullback-Leibler (KL) divergence (Fig 5b). The KL divergence between the output layer unit activities and the sampling distribution associated with the left target decreases as the total logLR supporting the left target gets larger and grows when the total logLR is smaller. The opposite trend is observed on the KL divergence between the output layer unit activities and the sampling distribution associated with the right target. These results suggest the activities of the output layer shape units encode the probability distribution of the next shape based on the current accumulated evidence.

### Task 2: Multi-sensory integration task

So far, we have shown that the network is able to perform the probabilistic reasoning task. The decision making process in the task may also be interpreted as Bayesian inference in which each shape updates the prior information. To more directly illustrate whether the network performs Bayesian inference, we test the model with a multi-sensory integration task adapted from Gu et al., 2008 (Gu et al., 2008) and compare it with the animals’ behavior. In this task, the subject has to discriminate whether the heading direction is toward the left or the right basing on either the visual input, the vestibular input, or both. Gu and colleagues showed that the animals were able to use information combined from different modalities optimally and reached a threshold that matched predictions based on Bayesian inference.

To model this task, we assume that the input layer units have Gaussian heading-direction tuning curves with Poisson noise. They are divided into two groups of 8 units, corresponding to the visual and vestibular inputs, respectively. Critically, the network is trained with trials in which inputs are from only one single sensory modality and tested with trials in which inputs are still limited to single modalities and trials in which inputs from both modalities are available. We find that the network can generalize the training to bi-modal inputs. It combines the information from visual and vestibular inputs and achieves a higher accuracy without further training (Fig 6a). We fit the network model’s choices in both the uni-modal and bi-modal conditions with cumulative Gaussian functions and define network’s performance thresholds as the standard deviation of the best-fitting function. The threshold under the combined condition is significantly smaller than the thresholds under either single modality conditions (Fig 6b). Importantly, it matches the predictions of Bayesian inference. These results suggest that the network model is able to perform Bayesian inference without being explicitly trained to do so.

**Figure 6.**
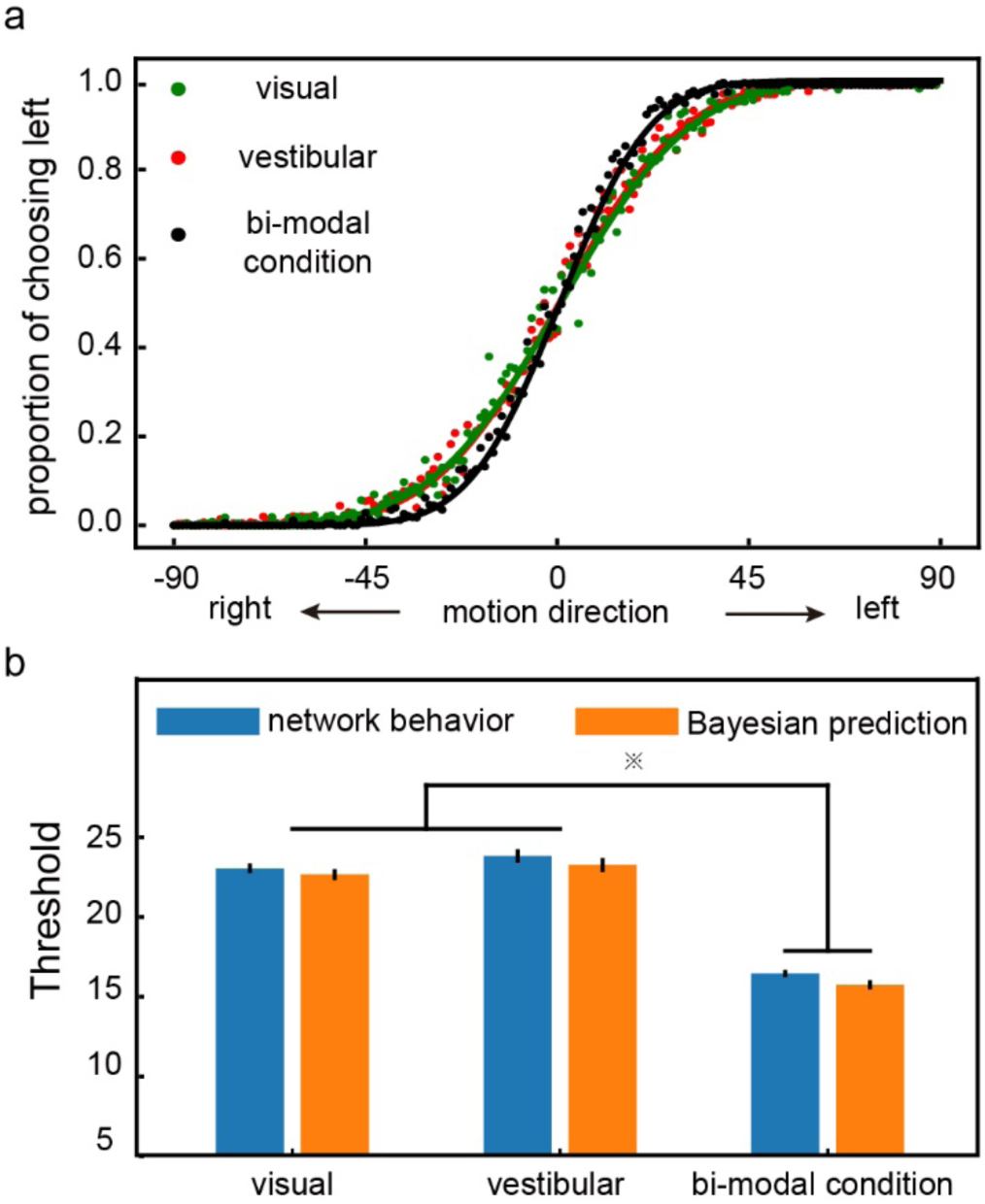
**a**. The psychometric curve. The model is first trained with the single modal conditions and then tested with both the single modal (green: visual, orange: vestibular) and the bi-modal (black) conditions. Each data point represents the proportion of the left choice at a given motion direction condition. The model shows a steeper psychometric curve for the bi-modal condition, indicating a better performance. **b**. The performance thresholds. The blue bars are the model’s thresholds under the single modal and the bi-modal conditions, compared against the thresholds calculated with the optimal Bayesian inference (orange). The threshold under the bi-modal condition is significantly smaller than the thresholds under either single modal condition. The differences between the thresholds of the network and the thresholds calculated with Bayesian inference are not significant. (two-tailed t-test with Bonferroni correction, p value threshold = 0.05)

### Task 3: Confidence / Post-decision wagering task

It has been argued that the same neurons that represent the decision variable during decision making may also support meta-cognition such as confidence. We test how our model performs a post-decision wagering task (Kiani & Shadlen, 2009). In this task, the model needs to make decisions about the movement direction of a random dot motion stimulus. The task difficulty is controlled by the proportion of dots moving coherently and the duration of the random dot stimulus. A reward is delivered if the decision matches the direction of the coherently moving dots. In half of the trials, a third target (sure target) appears after the motion viewing period. Choosing the sure target leads to a smaller but certain reward.

We set up the appropriate inputs and output units for our network model to learn the task. There are 10 input units for the motion stimulus, among which 5 prefer the leftward and the other 5 prefer the rightward motion. Their activities represent the motion strength of the random dots. Independent noises are added to the activities. Two additional input units indicate the left and the right choice target. Finally, there is an input unit that indicates the presence of the sure target. Again, the output units are a mirror copy of the input units, and the activities of the output units corresponding to the fixations on the choice targets and the sure target are used for generating the decisions.

The network exhibits a similar choice pattern to the monkeys’ behavior (Kiani & Shadlen, 2009). When the motion strength is weaker and the stimulus duration is shorter, the task difficulty is higher and the network chooses the sure target more frequently (Fig 7a). In addition, because the model may opt out for the sure target when its confidence about the motion direction is low, the overall accuracy becomes higher in trials in which the sure target is available but the model chooses to indicate the motion direction than in trials without the sure target (Fig 7b). The model’s behavior is consistent with the monkeys’ strategy.

**Figure 7.**
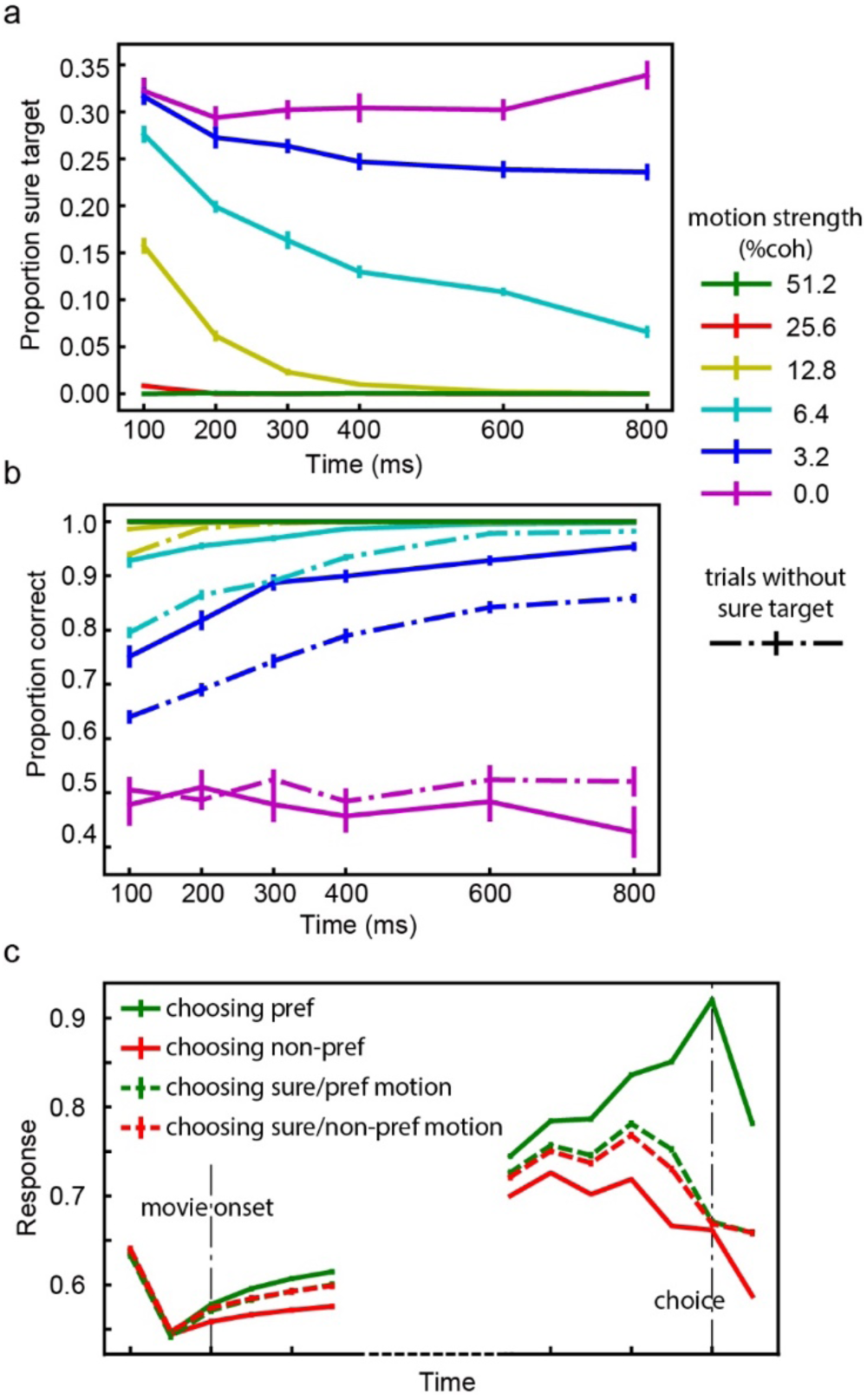
**a**. The proportion of trials in which the model chooses the sure target. The color indicates the motion strength. The frequency of the sure target choices decreases with the motion strength and the duration of motion viewing. b. The accuracy. Solid lines are the trials with the sure target and dashed lines are the trials without the sure target. The correct rate is higher when the sure target is provided. The error bars in panels a and b are S.E. across runs. c. Activities of choice-selective units. The responses are aligned to the movie onset (left) and the choice (right). The color of each line denotes the choice and motion direction. The dashed lines are trials in which the model chooses the sure targets. The units have intermediate activity level in trials that the sure target is chosen. The error bars are S.E. across units.

We further look at the response patterns of the units in the hidden layer. In particular, we want to study the units that show choice selectivity during the delay period before the choice. This selection criterium is similar to what was employed in the experimental study (Kiani and Shadlen, 2009, see Methods). The responses of these units are modulated during the motion viewing period (Fig 7c). More importantly, in the trials when the network chooses the sure target, their responses reach an intermediate level between the activity levels in trials when the targets of the preferred and non-preferred directions are chosen. These units behave like the neurons recorded from the LIP area and may provide a basis for the post-decision wagering (Kiani & Shadlen, 2009).

### Task 4: Two step task

In the last experiment, we extend our model to a learning task. Learning requires the model to incorporate trial history information for future decisions. By piecing events across multiple trials together, our network model should be able to infer contingencies across trials and achieve adaptive behavior. Notably, because the sensory, motor, and reward events are all included in our model, it is straightforward to achieve across-trial learning with our model.

To demonstrate that the network adapts well in changing environment, we use an adapted version of the origin two-step task (Akam, Costa, & Dayan, 2015; Daw et al., 2011). In this task, the agent has to choose between two options A1 and A2, each leading to one of the two intermediate outcomes, B1 and B2, with different but fixed transition probabilities. In particular, A1 is more likely followed by B1 (common) than B2 (rare) and A2 is more likely to be followed by B2 (common) than B1 (rare). Finally, B1 and B2 are each associated with a probabilistic reward (0.8 vs 0.2). The reward contingencies of B1 and B2 are reversed across blocks. Each block consists of 50 trials. The task requires the agent to learn that the intermediate states B1 and B2, instead of the agent’s own choice of A1 or A2, are actually the ones that determine the reward.

The training dataset consists of trials with randomly assigned choices. This is to reflect that the actual learning by human subject starts from rather random choices. During the training, only the events in the rewarded trials are actually trained. This is because the goal of the training is to learn the sequences more likely leading to reward. For the network to learn the event contingencies across trials, the state vector *h_t_* is not re-initialized at the beginning of each trial and the events in the previous trial are included in the training procedure. The loss function is calculated based only on the current rewarded trial, but the errors are back-propagated to the previous trial and the network connections are updated accordingly. In the testing sessions, the network connections are not further updated.

After the training, the network learns to make decisions based on the events both in the current and in the previous trials. Immediately after a block change, the network performs poorly because of the reversed reward contingency. Its performance then recovers gradually (Fig 8a). To exclude the possibility that the network simply learns to reverse its choice every 50 trials, we train the network with 50-trial blocks and test it with a different block size and observe similar results (Supp fig 2).

**Figure 8.**
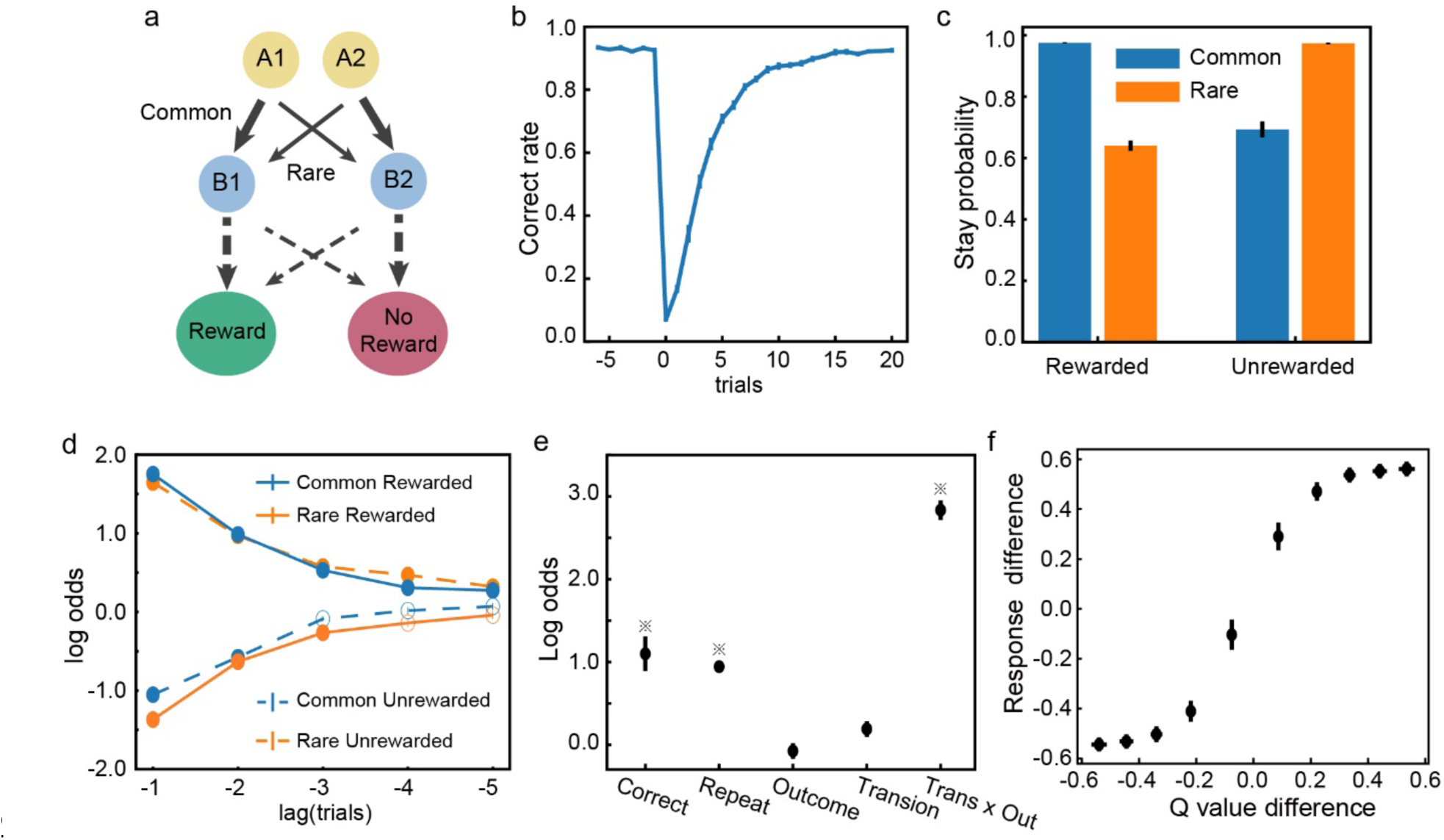
**a**. The two-step task. The thick and thin lines denote the common and rare transitions, respectively. The contingencies indicated by the dashed lines are reversed across blocks. **b**. The switching behavior. Trials are aligned to the block switch (trial 0). The performance first drops to below the chance level but then gradually recovers. **c**. The probability of repeating the previous choice. The stay probabilities of the subsequent trials are higher for the CR and the RU trials than the RR and the CU trials. **d**. Trial history effects. The choice in the current trial is affected by the trial types in the previous trials. Solid dots indicate significant effect (Bonferroni correction, p < 0.05). **e**. Factors affecting the choices. ※ indicates significance (p<0.01). **f**. The response difference between two choice output units is correlated with the difference between the estimated Q-value of the two actions.

To further investigate the network’s adaptive behavior, we use a factorial analysis introduced by Daw and colleagues (Daw et al., 2011) to look at how different factors of the task affect the choice. We sort the trials based on the intermediate outcomes and the reward outcomes into four groups: common-rewarded (CR), common-unrewarded (CU), rare-rewarded (RR) and rare-unrewarded (RU). Then we calculate the probability of the network repeating the previous choice in the next trial. The probability is higher after the CR and RU trials than after the CU and RR condition (Fig. 8b). The results indicate that the task structure information is used for decisions, which was interpreted as evidence for model-based decision making (Daw et al., 2011). Interestingly, even though the error signals are only propagated one trial back during the training period, the history influence on the network’s choice extends to many trials back (Fig 8c).

Two more analyses corroborate this conclusion. We first use a logistic regression to look at how a variety of factors affect network’s choices (Fig 8d). The result suggests that the interaction term between the intermediate outcome and the reward outcome (*Trans x Out*) is the largest factor affecting the choices. We further fit the network’s behavior with a mixture of the model-free and model-based agent (Akam et al., 2015; Daw et al., 2011). The higher weight for the model-based strategy (Table 1) indicates the network’s choice behavior is closer to model-based.

**Table 1.** Fitted parameters. The S.E. across runs is indicated. The weight for the model-based algorithm, wmodel_based, is significantly larger than 0.5 (p<0.001), indicating a strong contribution of the model-based behavior.

| $w_{model\_based}$ | $\alpha_1$ | $\lambda$ | $\alpha_2$ | $\beta$ | $p$ |
| --- | --- | --- | --- | --- | --- |
| $0.87 \pm 0.04$ | $0.75 \pm 0.05$ | $0.68 \pm 0.05$ | $0.47 \pm 0.01$ | $9.49 \pm 1.16$ | $0.77 \pm 0.08$ |

Although the network is not trained explicitly to calculate and compare the value associated with each state, it is curious whether the network activities encode the Q-value estimated with RL. Based the RL model fitting above, we estimate the choices’ Q-value in each trial. When we plot the estimated Q-value difference between the two choices, it shows a clear correlation with the activity difference between the two choice output units (Figure 8e).

## Discussion

Here, we present a neural network model for flexible and adaptive decision making. By training essentially the same network model to perform four exemplary tasks, we show that the network model not only can solve complicated decision-making problems, perform Bayesian inference, and support meta-cognition, but also can use past trial history to adapt its choices in a volatile environment. The units in the network exhibit signature behaviors of the decision-related neurons previously described experimentally, encoding the accumulative evidence and urgency signal. We also suggest a possible mechanism of the speed and accuracy trade-off. The network model provides a unified solution to questions that have been investigated previously with different modeling approaches.

### Drift-diffusion model and Attractor network models

The probabilistic reasoning task and the post-decision wagering task have been studied with the drift-diffusion model (DDM). In the DDM, the decision-making process is modeled as a diffusion process in which the evidence biases the drift rate. The behavior of the DDM can be accounted for by our model. This is interesting because, unlike the DDM and its variants, our model does not explicitly model the decision as a competition between alternatives in which evidence is accumulated. Yet by training the model to learn the statistical contingencies between events, the model exhibits behavior that fits the DDM well.

Our modeling effort is a distinct approach from the previous work of attractor neural network models (X.-J. Wang, 2002; Wong & Wang, 2006). Like the DDM, the attractor network models treat the decision-making process as a competition. The competition in attractor networks is between two groups of neurons, each receiving sensory inputs supporting its corresponding option. The sensory inputs drive the network states into attractor states that represent the final choice. This type of attractor network in its simplest form is obviously not a general solution for more complex decision-making tasks, for example, when the decisions require multiple steps as in the two step tasks, or when the contingency between the sensory inputs and the reward is volatile.

Even if we limit our discussions to the kind of decision making problem that has been studied with the attractor network model, the current approach provides several new insights. First, our model describes a potential mechanism for the speed-accuracy tradeoff in decision making distinct from the attractor models. In our model, there are a group of units that contribute to reaction time causally. Through the modulation of these units, one can achieve speed-accuracy tradeoff. In contrast, the speed-accuracy tradeoff in attractor network models is achieved through the adjustment of the balance between excitatory and inhibitory connections, which effectively modulate how fast the network may settle into an attractor state (Lo, Wang, & Wang, 2015). Future experiments can be designed to distinguish these two scenarios. Second, our model’s decisions do not require the network to converge into attractor states. This allows the model to perform the post-decision wagering task without additional modules. The network’s state, i.e. the units’ activity level, not only reflects the decision outcome, but also may be used to estimate confidence. It was found to be necessary to introduce an extra layer to model such a task with a attractor network (Insabato, Pannunzi, Rolls, & Deco, 2010). Finally, our model does not require its inputs to reflect evidence strength, which is essential for an attractor network. Therefore, it is able to perform the probability reasoning task in which the contingencies between the sensory stimuli and the outcomes are arbitrary. The input flexibility allows our model to be a more general solution than the attractor network models.

### Bayesian Inference

The computation that the network carries out is essentially Bayesian inference. During decision making, our network provides a representation of the probability distribution of the sensory events, which are updated over the time as new inputs arrive. Unlike the probabilistic population coding theory proposed previously (Ma, Beck, Latham, & Pouget, 2006), our model does not depend on assumptions of the distributions of the input or the nature of the contingency. The population response of the units in our model encode the entire distribution. This is a more general solution than the probabilistic population coding theory. Another recent model also aims to be a general solution for probabilistic computing (Orhan & Ma, 2017). In comparison, our model emphasizes the temporal integration of information. The outputs of our model may be directly mapped to the probability distribution, which may serve as a useful top-down signal that can be sent back to the sensory cortex.

### Reinforcement learning

By learning statistical relationships between sensory, action, and reward events across trials, our model can exhibit an adaptive behavior, which has been mostly modeled in the field with the reinforcement learning framework.

An interesting conceptual difference between our model and RL is the role of reward. The reward plays a central role in RL, in which the learning is aimed at minimizing the reward prediction error and maximizing the total future reward. The discrepancy between the actual and the expected rewards is used to update the value of states. In contrast, our model treats the reward events as part of the event sequences just as the sensory and action events. The learning in our model is driven by a teaching signal that includes sensory, action, and reward events. This signal could come from the dopamine system in the brain. The dopamine neurons have been indicated to signal reward prediction error and used as evidence for the existence of a reinforcement learning system in the brain (Schultz et al., 1997). Yet, recent findings have started to reveal a wider role of dopamine neurons in signaling not just reward prediction error, but also predictions of sensory stimuli or actions (Engelhard et al., 2018; Jin & Costa, 2010; Wassum, Ostlund, & Maidment, 2012; Wood, Simon, Koerner, Kass, & Moghaddam, 2017). These findings fit well with the current model in which the prediction errors need to be calculated for not only the rewards, but also the sensory and action events. Future experiments may be designed to test whether the dopamine system indeed fit this role.

Interestingly, our simulation results of the two-step task, along with several previous modeling studies (J. X. Wang et al., 2018; Zhang et al., 2018), show that online updating of synaptic weights is not necessary for adaptive behavior. Once the network is trained and learns the contingency between the events across trials, it can generate adaptive behavior based on the internal network dynamics, which encodes the past trial sequences, without further adjusting its internal connection weights. The network may appear to be learning and its behavior can be well explained by the reinforcement learning framework, yet no actual learning has to happen at the level of neuronal connections.

### Training

To train the model, it is necessary to feed the model with appropriate sequences of sensory, action, and reward events. In the real brain, this means there has to be a separate mechanism to generate appropriate responses at appropriate times and form these sequences in the first place for the brain to learn. One solution is to start from the responses based on animals’ innate responses or other established stimulus-action associations relevant to the task. These responses should contain a certain degree of variations that lead to different responses and reward outcomes (Neuringer, 2002). For example, when we train monkeys to perform complicated behavior task such as the probabilistic reasoning task, we start from the most basic delayed saccade task. Monkeys have an innate tendency of making saccades toward newly appearing visual stimuli at variable delays. By selectively rewarding saccades with longer delays, we can train the monkeys to hold the fixation for an extended period before the saccade. More components of the task can be introduced into the task gradually this way. Similar strategy can be used to train our model. Starting with simple tasks that the network is capable of doing, even if only occasionally, we can selectively reinforce the behavior that leads to reward and use the event sequence leading to the reward for the next stage training, until we reach to the final version of the task.

### Basal Ganglia

It remains an interesting speculation which brain structures may carry out the computation as our network model does. Areas such as the prefrontal cortex, the posterior parietal cortex, the cerebellum, the hippocampus, and the basal ganglia receive a wide range of sensory, motor, and reward inputs and are theoretically possible to carry out the computations demonstrated here. Indeed, many studies of these brain structures find neurons that have similar response patterns as the units in our model.

Among these brain structures, the basal ganglia are particular interesting. It has been long noticed that the key feature of GRU networks, which is the gating mechanism that controls the maintenance of old information and the updating with new incoming information, resembles the anatomical circuitry in the basal ganglia (Frank, Loughry, & O’Reilly, 2001; O’Reilly & Frank, 2006). The basal ganglia receive broad input from all over the cortex, including the sensory cortices and motor cortices (Engelhard et al., 2018), which comprises all the necessary information for learning sequences.

Our model may be applied to the current basal ganglia research in several distinct directions, which have been largely studied separately.

First of all, our modeling results explain previous studies of the basal ganglia’s role in decision making. For example, Ding and Gold found caudate neurons represented accumulated information in a random dot motion discrimination task (Ding & Gold, 2010, 2012). Cisek and colleagues observed a group of neurons in the globus pallidus showed activity patterns similar to the urgency signal (Thura & Cisek, 2017). In rodent experiments, it has been reported lesions in the striatum led to deficits in evidence integration and produced a bias in decision making (L. Wang, Rangarajan, Gerfen, & Krauzlis, 2018; Yartsev, Hanks, Yoon, & Brody, 2018). Our model may account for the results from these studies.

Second, our model, which is trained to learn sequences, naturally explains the basal ganglia’s role in performing sequential actions and in procedure memory (Geddes, Li, & Jin, 2018; Jin & Costa, 2010; Wood et al., 2017). It is conceivable that simply setting the input and output units according to the task requirement, we may train the model to perform sequential actions.

Third, the basal ganglia have also been indicated to play a central role in habitual and goal-directed behavior, both are highly relevant to our model (Graybiel, 1995; Graybiel & Grafton, 2015). On the one hand, by design, our model is capable of generating habitual behavior after the training. The model simply follows a set sequence of events and carries out actions, which may be interpreted as habitual responses, even in the case that the actions have to be determined with the preceding sensory inputs in a complicated manner. On the other hand, our model may also contribute to goal-directed behavior. A key component of goal-directed behavior is the ability of assessing which actions may lead to a desired goal. Such assessment depends on the capability of making predictions of the actions’ consequences, which is exactly what our model can do.

Our model may therefore help us to link the existing studies of basal ganglia from distinct perspectives to form a coherent computational theory of the basal ganglia’s role. Future work should incorporate more details of the basal ganglia circuitry into the network structure.

### Further investigations

The current results focus on the similarities between the model and the behavior and physiological findings from the previous experiments. It would be interesting to further explore how tweaking components and parameters of the GRU network would affect these results. This knowledge should motivate the experimental investigation of the relevant neural circuitry.

We also have not investigated extensively how the particular loss function we use to train the network in the current study, which treats all events equally and is flat over time, can be improved. One possibility is to normalize the loss function within each domain of sensory, motor, and reward, and use an appropriate weight for each domain to reach a suitable balance. Another possibility is to weigh loss functions differently across the time, so that events remote from the reward in the sequence exerts less influence on learning. Our preliminary explorations in these directions have yielded similar results (not shown), but more detailed investigations may reveal a loss function that may create a network that models the brain better and provides us with further insights.

Our current investigation focuses on the hidden layer, which is where the learning and the decision making occurs. Conceptually, this is similar to the generative model in the predictive coding framework (Friston, 2005; Rao & Ballard, 1999). The output layer in the current study only serves a limited purpose for the verification of the network’s performance. One may extend the current work and construct a circuitry in which the model’s outputs are sent to a sensory module and modulate the sensory processing. The sensory module sends inputs back to the model and forms a loop. This way, we can create a complete model that may be further used to understand the neural circuitry in both the sensory and the motor parts of the brain.

Finally, GRU networks have been shown to be able to solve problems at much larger scales, for example, the natural language processing (Chung et al., 2014). Limited by the availability of the existing experimental studies, we have only tested the network with small-scale toy problems. As a result, we use only a small network with 128 units. Natural language processing demands the learning of complex contingency structures between elements of language contained in a sequence that can be very long. Although at a much more complex level, this is essentially the same learning problem as what we study with our current model. By expanding our network model with more units and more layers, we should be able to model much more complex behavior. It is an interesting parallelism between the studies of neural network solutions for natural language processing, which develops neural networks to solve a complex computational problem that the human brain does, and the brain modeling studies, which uses network models to understand the computations in the brain. Our study is a beginning to bridge the two fields for the benefits of both.

## Acknowledgements

We thank Zhongqiao Lin, Chechang Nie, Yang Xie, Wenyi Zhang for their help in all phases of the study, and Shan Yu for providing comments and advice. The authors declare no competing financial or nonfinancial interests.

## Methods

### Task 1: Probabilistic reasoning task

#### Network

Our model is a network that contains three layers: an input layer, a hidden layer based on gated recurrent units, and an output layer (Fig 1a). The input layer contains units that carry the information about the sensory, motor, and reward events. (*N_IL_*=20).

There is a total of 14 sensory input units. 10 of them represent the 10 shapes in the task. There are 3 additional units indicating the presence of the eye movement targets, including the fixation point, the left target, and the right target. We also include a unit that indicates the absence of any visual stimuli.

There are 4 action input units that represent the efference copies of the motor commands: 1 for fixation on the fixation point, 1 for saccading to and then fixating on the left target, 1 for saccading to and then fixating on the right target, and 1 for saccading to other locations, which is considered as a fixation break and aborts a trial.

Finally, 2 reward input units are included: one for reward and one for the absence of reward.

The output layer includes also 20 units that mirror the inputs.

We set the time step to 100ms in the model training and testing. Our analyses and conclusions remain valid for smaller time steps.

#### Behavior Analysis

The parameters of the network are fixed during testing. We generate 5000 random shape sequences of 25 shapes long as the testing data and feed them into the network model. A shape sequence is stopped whenever the output units associated with the saccades are triggered. If the network does not make a response before all 25 shapes have been presented or makes a response at an inappropriate time, the trial is aborted (172.50 ± 3.73 or 3.45 ± 0.07 % trials in each run, excluded in further analyses).

#### Subjective weights (Fig 2b)

We perform a logistic regression to assess how each kind of shape affects the choice. The probability of choice is a function of the sum of leverages, *Q*, provided by each kind of shape:

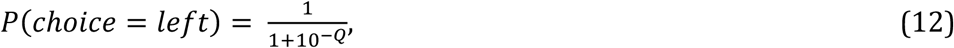

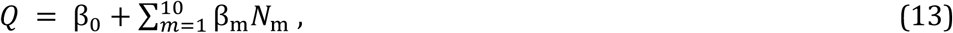

where *N_m_* represents the how many times shape *m* appears in a trial. β_0_ is the bias term, β_1∼10_ are the estimates of how much weight the network model assigns to shapes 1 to 10, and are termed as the subjective weights of evidence (Yang & Shadlen, 2007). Since the regressors are not independent to each other, here we use the ridge regression to minimize the variation of the estimation. The hyperparameter controls the trade-off between the cross-entropy loss and the L2-norm of the coefficients is selected through a ten-fold cross validation for each fitting.

#### Shape order (Fig 2d)

To test how the shape order affects the network’s choice, we perform a logistic regression on the trials with more than six epochs:

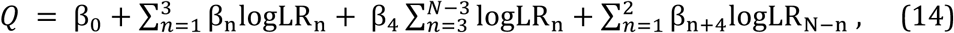

where logLR_n_ is the logLR of the shape in epoch *n* and *N* is the total number of epochs in a sequence. β_0_ is the bias term, β_1_, β_2_, β_3_ are the fitting coefficients of the shapes in the first three epochs, β_4_ the average effect for the middle epochs, and β_5_, β_6_ the second and the third epochs to the last. The regression is done without the final shape (*n*-th). This is because the last shape is almost always (>99% of trials) consistent with the choice. The fitting procedure is similar to what we use above to estimate the subjective weight of each shape.

#### Unit Selectivity (Fig 3ab)

We test whether each neuron’s activity in the hidden layer is modulated by the total logLR, absolute value of the total logLR, the urgency, and the choice outcome with a linear regression. We align the hidden unit state ℎ to the shape onset in each epoch and use a simple linear regression to characterize the unit’s selectivity:

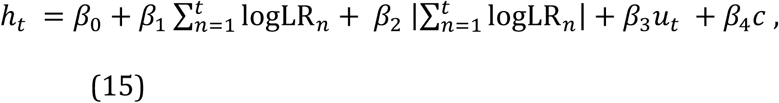

where *h_t_* is the unit’s response at time *t* aligned to the shape onset, 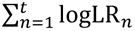 is the total logLR of the shapes that have been presented by time *t*, *u_t_* represents the urgency, which is quantified as the number of shape epochs, *c* represents the choice of the network, which is set to 1 when the left target is chosen, -1 when the right target is chosen, and 0 if fixation is still maintained by the end of the epoch. The regression is performed on every time step during the shape representation. We define a unit as selective to a variable if the variable shows a significant effect on the unit’s activity at every time step in an epoch. The significance is determined by two-tailed t-tests with Bonferroni correction.

#### Response Variability (Fig 3d)

We calculate a unit’s response variability as the standard deviation of the unit’s average response in each shape epoch across trials. Fig 3d plots the average response variability for all units that are selective to the total logLR in all 20 simulation runs, using only the trials with more than 5 shapes.

#### Speed-accuracy tradeoff (Fig 4d)

To test how *when* units affect the speed-accuracy tradeoff, we suppress the outputs of the +*when*/-*when* units that are selected with variable criteria. For example, the criteria of the accumulative *I_when_* of the selected +*when* units are set to 10%, 20%, 30%, 40%, and 50% out of the summed *I_when_* of all units with positive *I_when_*. At each criterium, the number of units manipulated is not linearly scaled, but their total contribution to the behavior, measured by their summed *I_when_*, is scaled linearly.

#### Network analysis (Fig 4cd)

Geodesic distance and max flow are used to quantify how much information may be transferred between two nodes in a graph. We use weight matrix *U_h_*, which represents the connection strengths between units in the hidden layer, to construct a weighted directed graph. Each unit in the hidden layer corresponds to a node in the graph. Connections with the highest 30% of the absolute value of the connection strength are turned into the edges in the graph. The results hold if we keep the highest 10% or 50%. The weight and capacity of each edge are defined as the absolute value of the original connection strength.

We define the geodesic distance from node A to target B as the minimum summed inverse of the weights of the edges from node A to B (Newman, 2001). The geodesic distances between any pairs are calculated with Dijkstra’s algorithm (Ahuja, Magnanti, & Orlin, 1993). The distance between the pairs not connected is defined as infinite. To account for these pairs, we compare the mean of the inverse of geodesic distance across all unit pairs in each group.

### Task 2: Multi-sensory integration task

#### Input units

There are two groups of input units representing the sensory stimuli, corresponding to the visual and the vestibular inputs. Each group has 8 units with Gaussian tuning curves with different preferred directions *D_pref_* at -90°, -64.3°, -38.6°, -12.9°, 12.9°, 38.6°, 64.3°, and 90°, respectively. The tuning curve *f*(*D*) is described as following:

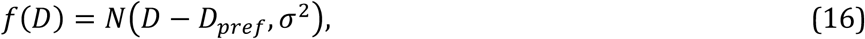

where *D* is the direction of the motion stimulus in the range between -90° and 90° and *σ*=90° is the standard deviation of the Gaussian function. The observed spike count *s* has a Poisson distribution:

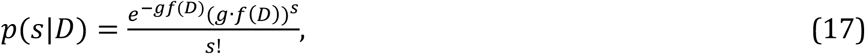

where *g*=300 is a constant for the gain. Finally, to limit the response of the input units to be smaller than 1, a sigmoid-like function is used to calculate the response *r*(*s*):

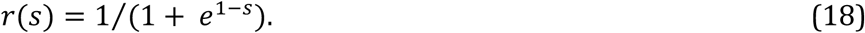

The exact choice of the function is not important. Under the bi-modal condition, the responses of both the visual and the vestibular input units are calculated with the heading direction *D*. Under the uni-modal condition, the responses of the units of the unavailable modality are set to 0.5 without noise.

#### Bayesian inference

We estimate the heading direction by calculating the discretized posterior probability of the possible motion directions;

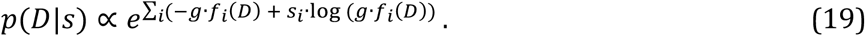

The left choice target is chosen if the probability of motion direction towards left, *p*(*D* < 0|*s*), is larger than the probability of motion direction towards right, *p*(*D* > 0|*s*). Otherwise, the right target is chosen.

In the bi-modal condition, we integrate the information from visual and vestibular system and calculate the posterior probability *p*(*D*|*s_vis_*, *s_vest_*), which is proportional to the product of the *p*(*D*|*s_vis_*) and *p*(*D*|*s_vest_*), given the independence between *s_vis_* and *s_vest_*.

All analyses are based on 20 simulation runs.

### Task 3: Confidence / Post-decision wagering task

#### Input units

The network has 22 input units. Ten input units are visual units that respond to motion stimuli. Five of them prefer the leftward motion and the other five prefer the rightward motion. Their internal activation, *s*, is linearly scaled with the coherence, and an independent Gaussian noise, *N*(0,*σ*^2^), is added to mimic the noise during the sensory processing, where σ=2.5. A sigmoid function is used as the activation function.

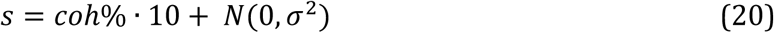

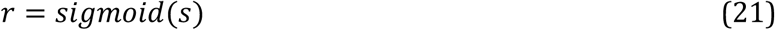

where the coherence of the moving dots, *coh*, is positive when the motion direction matches unit’s prefer direction and negative when not.

The momentary evidence *e_t_* is the response difference between the leftward-preferring units and the rightward-preferring units. The accumulated evidence *E* is then the sum of momentary evidence across time:

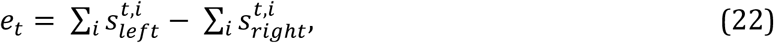

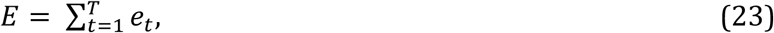

where 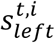 and 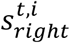 are the internal activities at time point *t* of the *i-*th unit preferring the leftward and the rightward motion, respectively. *T* is the duration of the motion stimulus and is randomly selected from group [1, 2, 3, 4, 6, 8] at equal probabilities. The final decision is based on the accumulated evidence *E*. The trial sequences used to train the model is generated as follows (Kiani & Shadlen, 2009). When the sure target is not available, the left target is chosen if *E* is larger than 0 and the right target is chosen if *E* is smaller than 0. When the sure target is given, the left and right choice targets are selected only if |*E*| is larger than a pre-defined threshold *θ*. Otherwise, the sure target is selected. Furthermore, the threshold *θ* increases linearly over time:

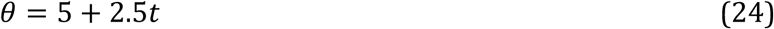

20 independent runs are simulated for testing the reproducibility.

#### Unit activity analysis

The analysis is based on choice selective units. These units are chosen based on their activities at the time point right before the choice in the trials without the sure target. The selectivity is determined by the two-tailed t-test with Bonferroni correction. The choice direction that induces a larger response is defined as a unit’s preferred direction (left: n = 46.75 ± 1.93, right: n = 42.45 ± 1.85).

### Two-step Task

#### Simulation

We use 20 trained session in our analysis. Each run contains 7.5*10^5^ training trials (15000 blocks of 50 trials) and 100 testing blocks.

#### Model fitting

The analysis was previously described (Akam et al., 2015; Daw et al., 2011; Zhang et al., 2018). Briefly, the model choice is fitted to a mixed model-free and model-based algorithm. For the model-free algorithm, the value of the chosen options and observed intermediate outcomes are updated by a temporal difference algorithm with two free parameters: the learning rate *α*1 and the eligibility *λ*, representing the proportion of the reward prediction error that is attributed to the first-stage chosen options A1 and A2. In the model-based algorithm, the value of the observed intermediate outcome is also learned with a temporal difference algorithm. The value of each option is the sum of the products of the transition probabilities and the values of the intermediate outcomes. An extra parameter, learning rate *α*2, is used for the update of the value of the intermediate outcomes in the model-based algorithm. The overall value of each option is the weighted average of its value calculated with the model-free and model-based algorithm (*w_model-free_*+ *w_model-based_*=1). Finally, the probability of choosing each option is a softmax function of the values of the two options:

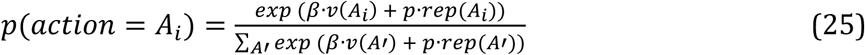

where inverse temperature parameter, *β*, controls the randomness of the choice. *rep*(*A_i_*) is set to 1 if action *A_i_* is chosen in the previous trial and the parameter *p* captures the tendency of repeating the previous trial. *v*(*A_i_*) is the value of option *A_i_*. Together, there are six free parameters and a maximum likelihood estimation algorithm is used for fitting.

#### Logistic regression

The analysis was described previously (Akam et al., 2015; Zhang et al., 2018). Briefly, five potential factors are tested with a logistic regression, which are *Correct*—a tendency to choose the choice with higher reward probability; *Repeat*—a tendency to repeat the choice no matter what the reward outcome is; *Outcome*—a tendency to repeat the rewarded choice in the previous trial; *Transition*—a tendency to repeat the choice when it leads to common intermediate outcome and switch when rare intermediate outcome is represented; *Trans x Out*–a tendency to repeat the same choice when the previous trial is CR or RU, and to switch the choice if the previous trial is CU or RR.

---

**Supplementary Figure 1.**
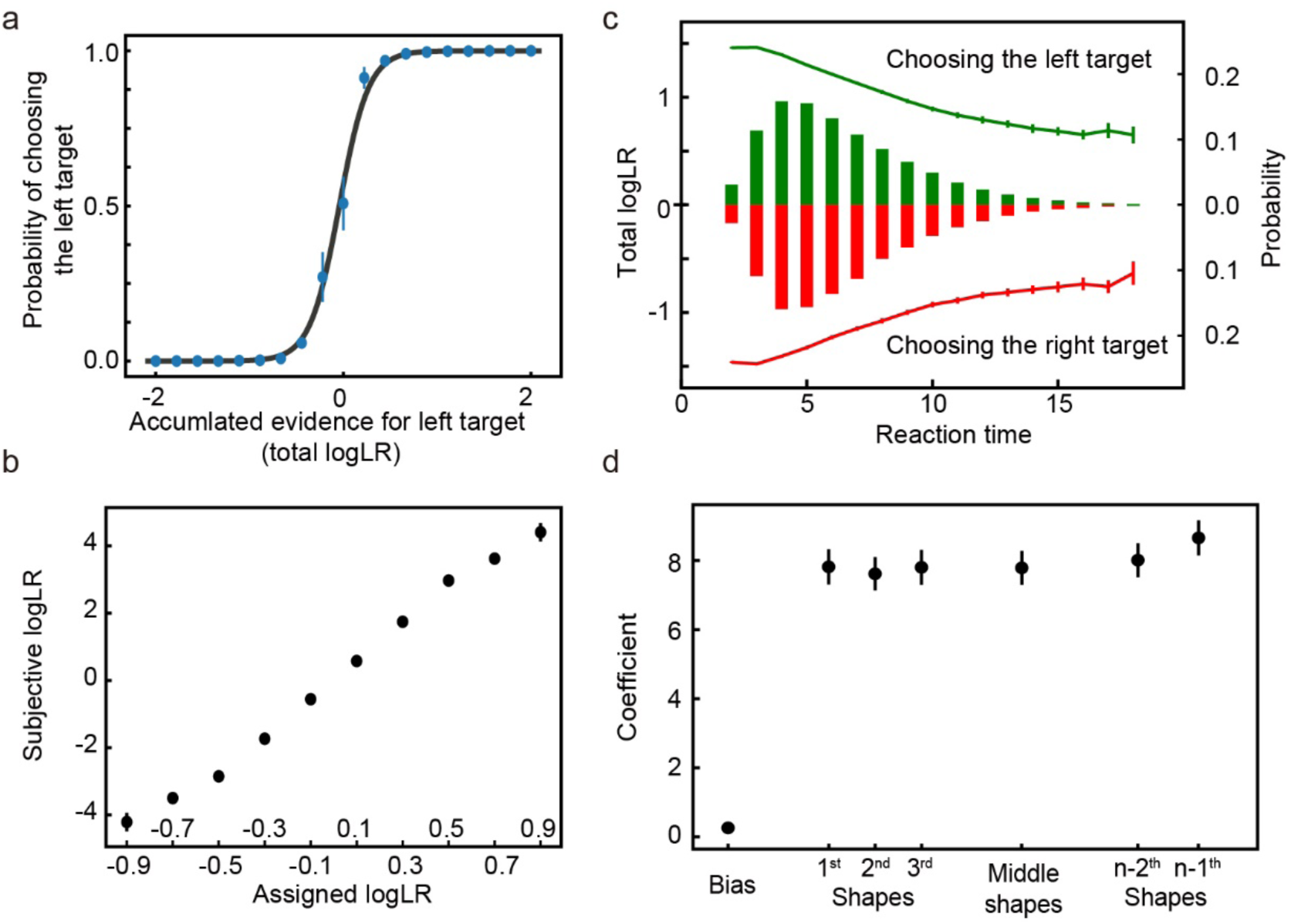
The performance of the network trained with a dataset that contains only 1000 unique sequences. Same format as in Figure 2.

**Supplementary Figure 2.**
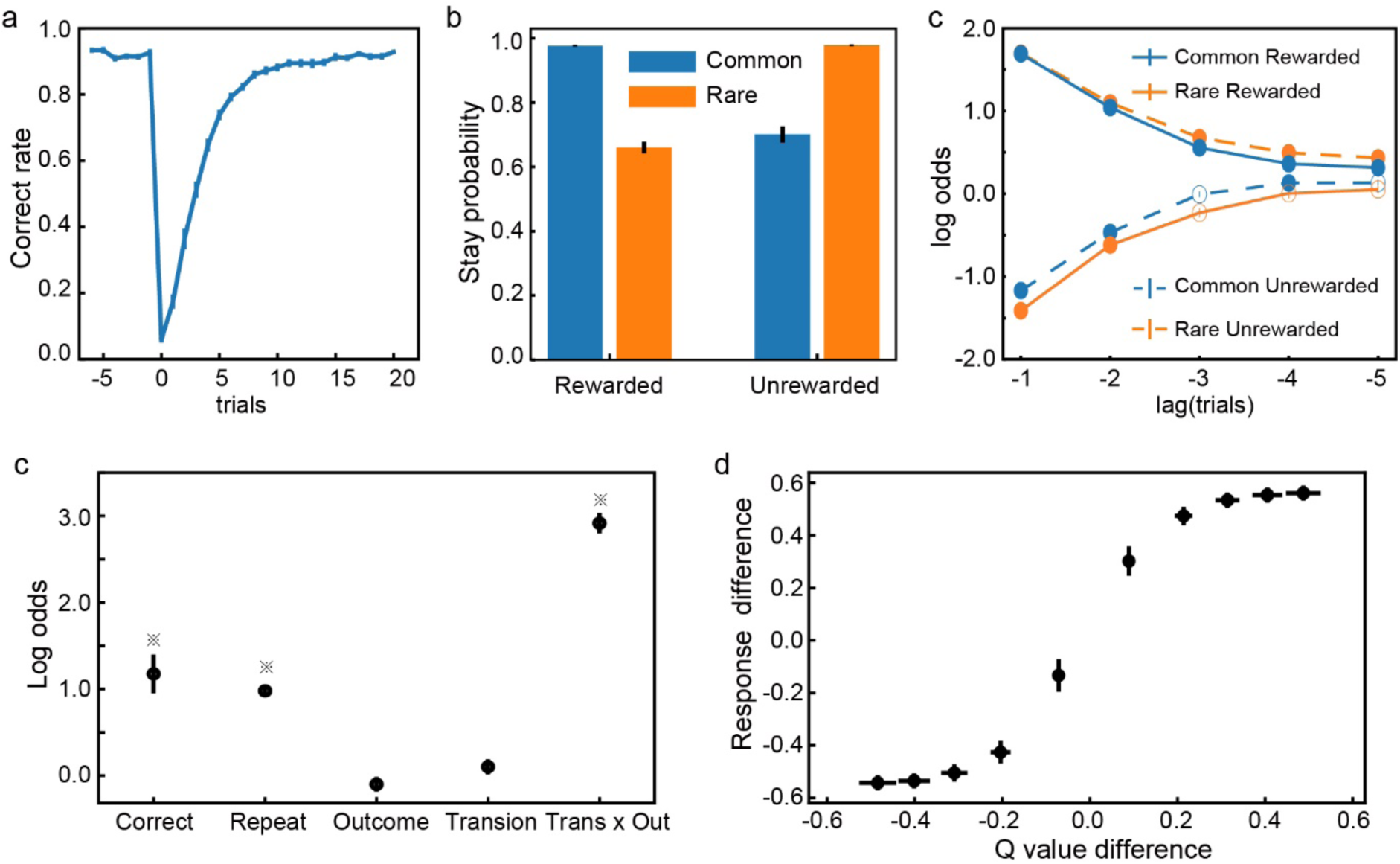
The network’s performance when the testing dataset’s block size is set to 70 trials. Same format as in Figure 8.

